# Age- and sex-dependent effects of DNA glycosylase Neil3 on amyloid pathology, adult neurogenesis, and memory in a mouse model of Alzheimer’s disease

**DOI:** 10.1101/2022.03.10.483733

**Authors:** Milena A. Egiazarian, Silje Strømstad, Teri Sakshaug, Ana B. Nunez-Nescolarde, Nicole Bethge, Magnar Bjørås, Katja Scheffler

## Abstract

Oxidative stress generating DNA damage has been shown to be a key characteristic in Alzheimer’s disease (AD). However, how it affects the pathogenesis of AD is not yet fully understood. Neil3 is a DNA glycosylase initiating repair of oxidative DNA base lesions and with a distinct expression pattern in proliferating cells. In brain, its function has been linked to hippocampal-dependent memory and to induction of neurogenesis after stroke and in prion disease. Here, we generated a novel AD mouse model deficient for Neil3 to study the impact of impaired oxidative base lesion repair on the pathogenesis of AD. Our results demonstrate an age-dependent decrease in amyloid-β (Aβ) plaque deposition in female Neil3-deficient AD mice, whereas no significant difference was observed in male mice. Furthermore, male but not female Neil3-deficient AD mice show reduced neural stem cell proliferation in the adult hippocampus and impaired working memory compared to controls. These effects seem to be independent of DNA repair as both sexes show increased level of oxidative base lesions in the hippocampus upon loss of Neil3. Thus, our findings suggest an age- and sex-dependent role of Neil3 in the progression of AD by altering cerebral Aβ accumulation and promoting adult hippocampal neurogenesis to maintain cognitive function.

## INTRODUCTION

Alzheimer’s disease (AD) is the most common cause of dementia with a complex pathogenesis that is not yet fully understood. Microscopically, it is characterized by extracellular amyloid plaques (Aβ), derived from amyloid precursor protein (APP), and intracellular neurofibrillary tangles (NFTs). Interestingly, AD is affecting more women than men, not only in prevalence, but also in severity [46]. Female patients exhibit a faster clinical decline than men [24, 38]. Mice models of AD show a greater amyloid load in female mice [7]. Thus, investigating sex differences is of importance to better understand AD pathogenesis, as well as improve diagnostics and treatment options.

Progressive memory impairment is associated with degeneration of the hippocampus, an area of the brain that is severely affected in AD [44, 45]. The dentate gyrus (DG) of the hippocampus is critical for learning and memory functions and harbors neural stem cells with the capacity to generate new neurons throughout life, a process termed neurogenesis. While the majority of neural stem cells remain quiescent in the adulthood, some contribute to the formation of new neurons by giving rise to transit-amplifying progenitor cells [30]. The decline of adult hippocampal neurogenesis found in patients with AD [44], might contribute to the impaired memory and cognitive dysfunction in AD patients [25, 63].

Oxidative stress is an integral part early in AD pathogenesis and has been shown to negatively affect adult neurogenesis [26, 34, 69]. Elevated levels of oxidative DNA damage have been found in post-mortem brains of AD patients, and in patients with mild cognitive impairment [41, 53]. Oxidative stress causing DNA damage is a result of an imbalance between free radicals and antioxidative detoxification. Aβ has been proposed to stimulate production of reactive oxygen species (ROS) by interacting directly with mitochondria, promoting oxidative stress and leakage of ROS [5, 20, 42, 60]. On the contrary, studies observed that oxidative damage anticipated Aβ accumulation and even triggered its production [2, 50].

Oxidative DNA damage is recognized by DNA glycosylases and repaired through the base excision repair (BER) pathway. The Neil-family glycosylases Neil1, Neil2 and Neil3 express overlapping substrate specificity, recognizing oxidized pyrimidines and purines [31]. Neil3 is expressed in specific cell cycle stages, peaking at S/G2 with a preference of repairing base lesions in single-stranded DNA (ssDNA) [21, 47]. Increased levels of 5OH-dC, a substrate of NEIL3, has been found in the parietal and temporal lobe of AD patients [39, 41, 43, 66]. In contrast to the other Neil-family members, Neil3 shows a cell-specific expression in proliferative cells of the testis, thymus and hematopoietic tissues, as well as the neurogenic niches of the brain [22, 40, 55, 57, 64]. Its function in brain has been associated with learning and memory, anxiety-like behavior, and neural regeneration after hypoxic-ischemic injury. Interestingly, Jalland et al. found a protective function of Neil3 during prion disease as a result of induced neurogenesis [28]. Altogether, these studies indicate a role for Neil3 in neurological diseases involving dysregulated neurogenesis such as AD.

This study aimed to investigate the impact of Neil3 on the progression of AD by generating and characterizing a novel Neil3-deficient AD mouse model.

## MATERIALS AND METHODS

### Mouse models

All mice used in our study were on a C57BL/6J background. APP/PS1 mice previously described by Radde et al. [51] and *Neil3^−/−^* mice previously described by Sejersted et al. [56] were used to generate APP/PS1x*Neil3^−/−^* mice. All experiments were performed with mice at 1 and 6 months of age. The mice were housed at ambient temperature on a 12:12 h light:dark cycle with free access to water and food.

### Immunohistochemical (IHC) and -fluorescent (IF) staining

Mice were transcardially perfused with PBS before brain dissection. For IF, mice were additionally injected with 100 mg/kg bodyweight BrdU (Sigma) 24 h prior to dissection. Brains were sliced into 4 μm thick coronal sections and embedded in paraffin. Sections were rehydrated in Xylol 2×5 min followed by ethanol 100% 2×3 min, 96% 1 min and 70% 1 min. For Aβ-plaque staining, the HRP/DAB-kit (Abcam) was used. Sections were first blocked in H_2_O_2_ for 10 min, and antigen retrieval was performed by incubating sections at 100°C citrate buffer (100 mM, pH 6.0) for 5 min (IHC) or 3 min (IF) under pressure. For IHC sections were blocked with Protein block for 10 min and incubated with primary antibodies against Aβ (6F3D Amyloid β, Thermo Fisher) diluted 1:100 in PBS + 0.5% goat serum + 0.5% BSA for 1 h at room temperature. Sections were thereafter incubated in Biotinylated Goat Anti-Polyvalent and Streptavidin Peroxidase for 10 min each and treated with DAB for 1-10 min. Sections were washed three times with washing buffer (PBS + 0.2% Tween 20) and incubated in hematoxylin (Sigma Aldrich) for 3 min, 98% ethanol + 2% NH_4_OH for 1 min, Eosin Y (Sigma Aldrich) for 15 s before dehydration followed by mounting (Entellan mounting medium, Sigma Aldrich). Sections were scanned by the Cellular and Molecular Imaging Core facility (CMIC, NTNU) using an OlympusVS120S5 with 20X objective, scanned on one Z-plane with low focus grid density. Plaque quantification was done using ImageJ. Color-presets were chosen to minimize background to count and measure number and size of Aβ-plaques. For the final quantification, the average of at least two sections per animal stained in independent runs were used.

For IF staining, sections were blocked in blocking solution (5% BSA + 5% goat serum in PBS) for 1 h and incubated with rat anti-bromodeoxyuridine (BrdU) 1:400 diluted in PBS + 0.5% BSA + 0.5% goat serum. Sections were washed in wash buffer (PBS + 0.2% Tween 20) and incubated with DAPI staining solution (1 μg/ml, Invitrogen) for 10 min before mounted in Vectashield mounting medium (Vector Laboratories). Microscopy was carried out using a Zeiss LSM 510 Meta confocal laser scanning microscope equipped with a 40X oil immersion lens, and quantifications were performed manually and blinded. For the final quantification, the average of at least two sections per animal stained in independent runs were used.

### Extraction of nucleic acids

Extraction of total DNA and RNA was performed using the Allprep DNA/RNA/Protein kit (Qiagen) according to the manufacturers protocol. Initially, 350 μL Buffer RLT + β-ME was added to the brain tissue and samples were homogenized with MagNA Lyser for 10 s at 5000 speed followed by centrifugation for 3 min at 16 400 rpm before proceeding according to the kits protocol. DNA and RNA concentrations were measured using NanoDrop (Thermo Fisher).

### Analysis of mRNA expression level

RNA from brain samples was reversely transcribed to cDNA with the High Capacity cDNA Reverse Transcription kit (Applied Biosystems) using T100 Thermal Cycler (Bio-Rad), programmed to run at 25°C for 10 min, 37°C for 120 min and 85°C for 5 min. The qPCR was performed using power SYBR Green PCR MM (Applied Biosystems) and run on a StepOnePlus Real-Time PCR System (Applied Biosystems). Primer sequences used for *Neil3* were GGGCAACATCATCAAAAATGAA forward and CTGTTTGTCTGATAGTTGACACACCTT reverse. Relative gene expression was calculated by using the ΔCt method and normalized to the expression of the housekeeping gene β-actin with primer sequences CTTGATAGTTCGCCATGGAT forward and GGTCACTTACCTGGTGCCTA reverse.

### RNA sequencing

Total RNA from whole brain of one male and one female APP/PS1 and APP/PS1x*Neil3^−/−^* mice, respectively per age group was sent to BGI Tech Solutions Co., Hong Kong, for RNA sequencing on an Illumina HiSeq4000. Unpaired reads were trimmed and filtered for quality. Quality of the trimmed reads were assessed using FastQC (v0.11.9) and FastQ-Screen (v.0.14.1). Reads were mapped to the GRCm38 transcriptome using Hisat2 and reads per gene were counted using the Rsubread package in R (3.6.3). EdgeR (3.32.1) was used to generate a log2 CPM matrix which was then exported to Perseus (v.1.6.14.0) to generate heatmaps. Male and female counts were separated, Z-scored by row and clustered using Euclidean hierarchical clustering. Gene lists from clusters were exported and analyzed using PantherDB for an overrepresentation analysis looking at Gene Ontology (GO) biological pathways.

### Quantification of oxidative DNA damage by mass spectrometry

DNA samples were digested by incubation with a mixture of nuclease P1 from Penicillium citrinum (Sigma), DNaseI (Roche) and ALP from E. coli (Sigma) in 10 mM ammonium acetate buffer pH 5.3, 5 mM MgCl_2_ and 1 mM CaCl_2_ for 30 min at 40ºC. The samples were methanol precipitated, supernatants were vacuum centrifuged at room temperature until dry, and dissolved in 50 μl of water for LC/MS/MS analysis. Quantification was performed with the use of an LC-20AD HPLC system (Shimadzu) coupled to an API 5000 triple quadrupole (ABSciex) operating in positive electrospray ionization mode. The chromatographic separation was performed with the use of an Ascentis Express C18 2.7 μm 150 × 2.1 mm i.d. column protected with an Ascentis Express Cartridge Guard Column (Supelco Analytical) with an Exp Titanium Hybrid Ferrule (Optimize Technologies Inc.). The mobile phase consisted of A (water, 0.1% formic acid) and B (methanol, 0.1% formic acid) solutions. The following conditions were employed for chromatography: For unmodified nucleosides – 0.13 mL/min flow, starting at 10% B for 0.1 min, ramping to 60% B over 2.4 min and re-equilibrating with 10% B for 4.5 min; For 5-OH(dC) - 0.14 mL/min flow, starting at 5% B for 0.1 min, ramping to 70% B over 2.7 min and re-equilibrating with 5% B for 5.2 min; For 8-oxo(dG) - 0.14 mL/min flow, starting at 5% B for 0.5 min, ramping to 45% B over 8 min and re-equilibrating with 5% B for 5.5 min. For mass spectrometry detection the multiple reaction monitoring (MRM) was implemented using the following mass transitions: 252.2/136.1 (dA), 228.2/112.1 (dC), 268.2/152.1 (dG), 243.2/127.0 (dT), 244.1/128 [5-OH(dC)], 284.1/168.1 [8-oxo(dG)].

### Human Aβ42 measurement by ELISA

Whole brain samples were homogenized in carbonate buffer (pH 11.5, 100 mM Na_2_CO_3_, 50 mM NaCl, protease inhibitor) using the MagNA Lyser (Roche) for 5 s at 5000 speed. Samples were centrifuged for 90 min at 16400 rpm. The supernatant was diluted in 8M guanidine buffer (pH 8, guanidine/HCl 8.2 M, Tris/HCl 83 mM) to obtain soluble Aβ42 fraction. For the insoluble Aβ42, the remaining pellets were resuspended in 5 M guanidine buffer (pH 8, guanidine/HCl 5M, Tris/HCl 50 mM) and shaken at room temperature for 3 h. Samples were centrifuged for 20 min at 16 400 rpm. Protein concentration was measured with NanoDrop (Thermo Fisher). Aβ42 concentrations were measured using the Human Aβ42 ELISA kit (Thermo Fisher) according to the manufacturer’s protocol. Absorbance was read at 450 nm with FLUOstar Omega (BMG Labtech).

### Behavioral analysis

The Elevated Zero Maze (EZM) assessing activity and anxiety behavior was performed on a circular runway 60 cm above the floor with four alternating open and closed areas. The mice were placed on the maze facing a closed area and allowed 5 min for free exploration of the apparatus. The T-Maze was used to measure working memory by assessing spontaneous alternation in a T-shaped arena according to the protocol described by Deacon and Rawlins [14]. A total of 6 training and test runs, distributed across two days with 1 h in between was performed per animal. During the training, mice were placed in the start arm and allowed to choose a goal arm within a maximum of 90 s. Mice were confined for 30 s in the chosen arm and then removed from the arena for 60 s. In the test run, mice were placed back into the start arm and allowed to choose between the two goal arms within a maximum of 90 s. A successful alternation was scored if mice choose to enter the previously non-visited goal arm in the training. During the behavioral tasks mice were monitored and analyzed by using ANY-maze video tracking system (Stoelting).

### Statistics

Statistical analyses were done with GraphPad Prism software version 8, using 2-way ANOVA with Tukey’s multiple comparisons test. P-values < 0.05 were considered significant (*p < 0.05, **p <0.01, *** p<0.001, ****p < 0.0001). Figures are shown with mean + standard deviation.

## RESULTS

### Reduced plaque deposition upon loss of Neil3 in female AD mice

We generated a novel APP/PS1x*Neil3^−/−^* mouse model to study how Neil3 affects AD pathology. The mice were viable and fertile, and showed no differences in weight compared to the APP/PS1 mice (Additional file 1: Fig. 1). To assess whether Neil3 has an impact on amyloidogenesis we first stained for Aβ plaques in the brain of the AD mice strains during disease progression. (Fig. 1A). Quantification of plaque numbers demonstrated an age-dependent increase from 1 to 6 months in all strains (Fig. 1B). Female APP/PS1 mice had higher plaque numbers at 6 months compared to its male counterparts. In contrast, Neil3-deficient AD mice showed no sex-specific change in Aβ deposition, thus female APP/PS1x*Neil3^−/−^* mice exhibited a significantly lower number of Aβ plaques compared to female APP/PS1. However, this finding was not reflected measuring Aβ42 concentrations in whole brain by ELISA (Additional file 2: Fig. 2). Although female APP/PS1 mice showed a tendency to higher guanidine-soluble (mostly fibrillar material) Aβ42 compared to male mice, no significant difference was found upon loss of Neil3 (Additional file 2: Fig. 2A). Similarly, buffer-soluble (mostly monomers and small oligomers) level of Aβ42 was comparable between the mouse strains at the age groups analyzed (Additional file 2: Fig. 2B). This suggests that Neil3 is involved in plaque pathology in a sex-dependent manner, but not in cerebral Aβ42 accumulation.

**Fig. 1.**
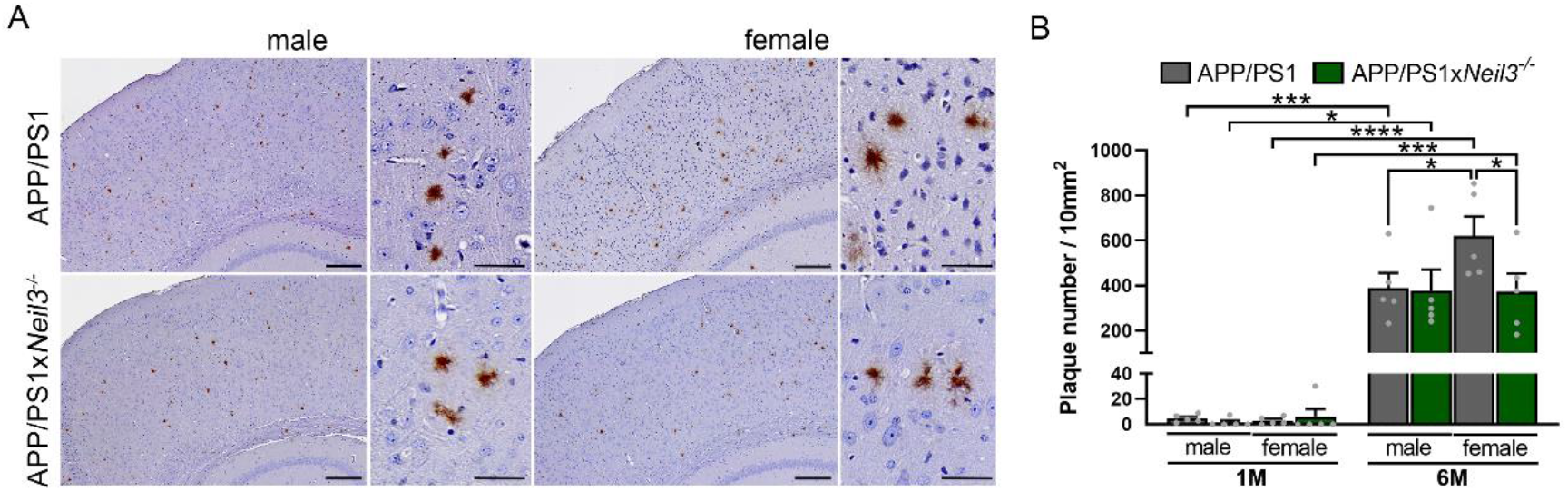
Loss of Neil3 decreases cerebral plaque deposition in female APP/PS1 mice with age. (A) Representative images of immunohistochemically stained amyloid-β plaques in the cortex and (B) quantification of plaque number in the brain of APP/PS1 and APP/PS1x*Neil3^−/−^* mice at 1 month (1M) and 6 months (6M) of age. Data are shown as mean + SD; n=5 per group. *p < 0.05, **p <0.01, *** p<0.001, ****p < 0.0001.

**Fig. 2.**
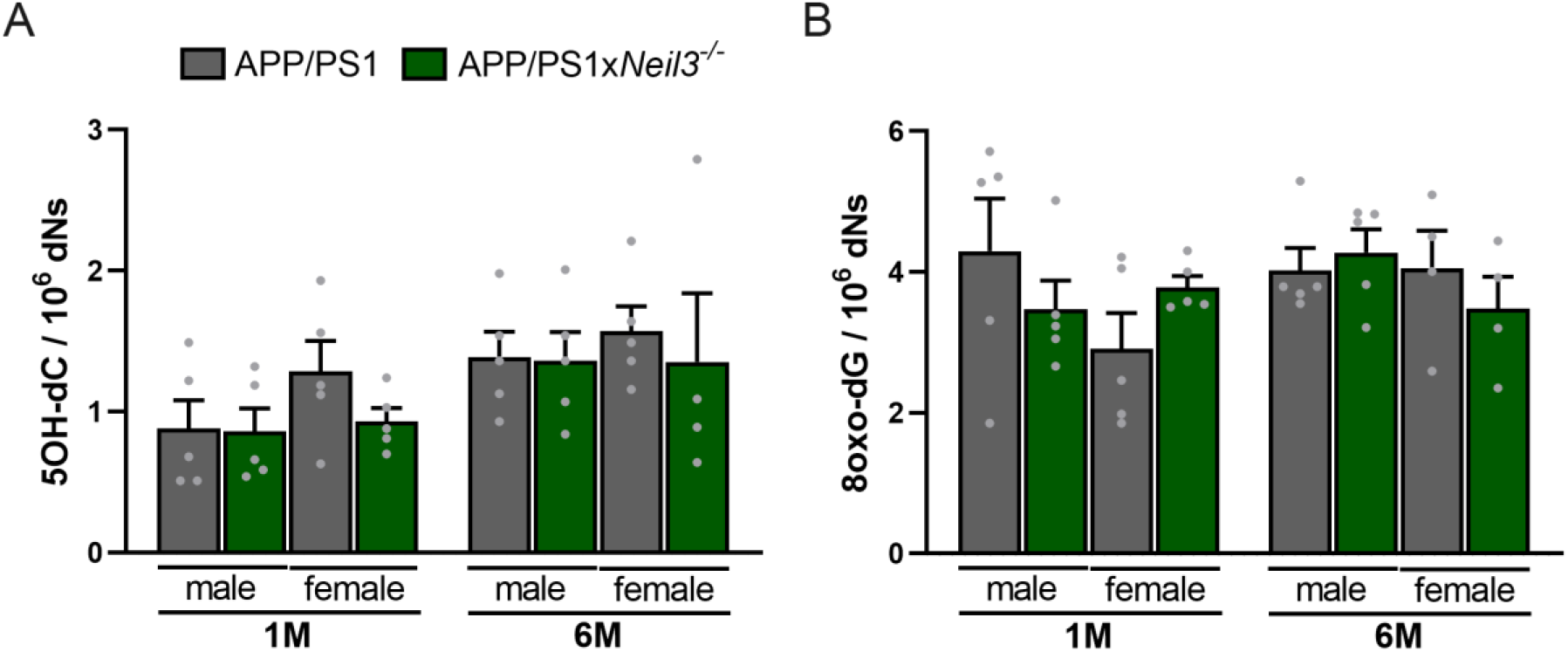
No differences in cerebral accumulation of oxidative base lesions in the Neil3-deficient AD mouse model. (A) Levels of oxidized base lesions 5-hydroxycytosine (5OH-dC) and (B) 8-oxoguanine (8oxo-dG) in brain samples of APP/PS1 and APP/PS1x*Neil3^−/−^* mice at 1 month (1M) and 6 months (6M) of age measured by using mass spectrometry. Data are shown as mean + SD; n=4-5 per group.

### No differences in levels of global oxidative DNA damage

Elevated levels of 5OH-dC, a known substrate of NEIL3 has been found in post-mortem brains of AD patients [18, 40, 41, 65, 66]; thus, we quantified the level by LC-MS in the AD mouse models (Fig. 2A). We found no differences in cerebral 5OH-dC levels neither between genotypes, nor age. In addition, levels of the common marker of DNA oxidation 8oxo-dG that was shown to accumulate during AD progression [18, 48, 58] were comparable between the mice strains and age groups (Fig. 2B). This indicates that Neil3 is dispensable for global oxidative DNA base damage repair in the brain of AD mice at the time points analyzed. Moreover, increased oxidative DNA damage seems not to be a hallmark in our AD mouse model until 6 months of age.

### Age-dependent impairment of working memory upon loss of Neil3 in male AD mice

As Neil3 was shown to affect anxiety, learning and memory in adult mice [52], we investigated whether Neil3 deficiency also affects behavior in the AD mouse model. The Elevated Zero Maze (EZM) was used to examine activity measured as distance traveled and anxiety levels measured by the time spent in the open arm. We found no major differences in activity and anxiety behavior between genotypes and age groups (Fig. 3A and B). Only Neil3-deficient male AD mice were significantly less active from 1 month to 6 months of age (Fig. 3A). The alternation rate as a measure of working memory was assessed in a T-Maze. Whereas AD progression from 1 month to 6 months did not affect working memory, loss of Neil3 led to a significant age-dependent decrease of working memory exclusively in male AD mice (Fig. 3C). In contrast, wildtype (WT) and *Neil3^−/−^* male mice demonstrated similar working memory at 6 months of age (Additional file 3: Fig. 3). This indicates that the effect of Neil3 on working memory is age- and sex-dependent and specific to AD.

**Fig. 3.**
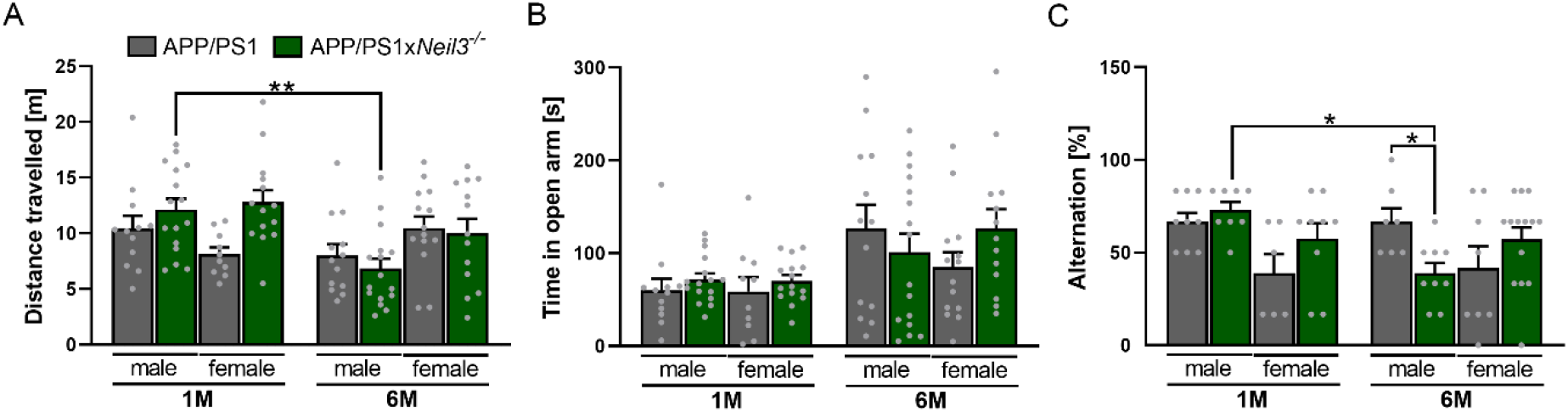
Behavioral testing reveals impaired working memory with age upon loss of Neil3 in male AD mice. (A) Distance travelled and (B) Time spent in the open arm assessed by using an elevated zero maze, and (C) Alternation rate assessed in a T-maze of APP/PS1 and APP/PS1x*Neil3^−/−^* mice at 1 month (1M) and 6 months (6M) of age. Data are shown as mean + SD; n=6-16 per group. *p < 0.05, **p <0.01, *** p<0.001, ****p < 0.0001.

### Impaired adult neurogenesis upon loss of Neil3 in male AD mice

Adult neurogenesis in the hippocampus has been shown to be important for cognitive functions [1, 15, 59] and Neil3 is expressed in stem cells of the hippocampus. To address whether impaired adult hippocampal neurogenesis may contribute to memory impairment in Neil3-deficient mice during AD progression, we used BrdU injection to stain proliferating cells in the DG (Fig. 4A). We observed significant fewer dividing cells in Neil3-deficient male AD mice at 1 month of age compared to male controls (Fig. 4B). Additionally, number of BrdU^+^ cells dropped drastically from 1 to 6 months of age in all AD mouse strains, suggesting an age-dependent decrease in proliferation capacity of DG cells during AD progression.

**Fig 4.**
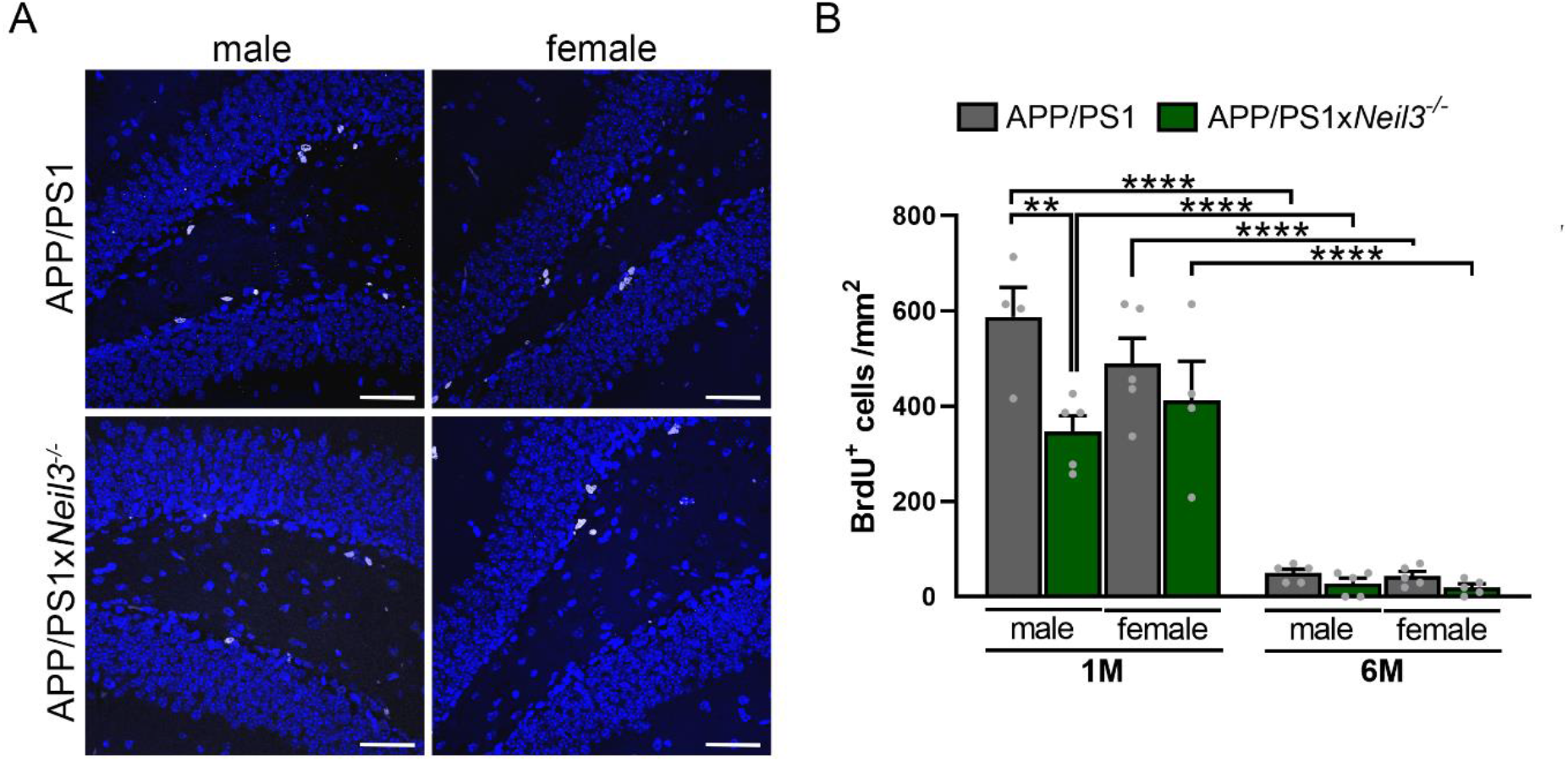
Lack of Neil3 leads to reduced stem cell proliferation in the dentate gyrus (DG) of young male AD mice. (A) Representative images of BrdU-stained cells (white) and DAPI-stained nuclei (blue) in the DG of 1 month old APP/PS1 and APP/PS1x*Neil3^−/−^* mice. Scale bar 50μm. (B) Quantification of BrdU-positive cells in the DG of APP/PS1 and APP/PS1x*Neil3^−/−^* mice at 1 month (1M) and 6 months (6M) of age. Data are shown as mean + SD; n=4-5 per group. *p < 0.05, **p <0.01, *** p<0.001, ****p < 0.0001.

### RNA sequencing reveals sex-specific differences in expression of genes involved in synaptic processes

To identify potential molecular pathways that could explain the effect of Neil3 in AD pathogenesis, we analyzed the transcriptome in the whole brain of the AD mouse strains at 1 month and 6 months of age by RNA sequencing. Hierarchical clustering with GO pathway analysis revealed sex-specific differences in AD-related processes (Additional File 4: Fig. 4). Male AD mice showed a significant enrichment of genes in inflammatory processes that increased in expression (cluster 1) and in postsynaptic processes that decreased in expression (cluster 10) from 1 month to 6 months of age (Fig. 4A). Interestingly in cluster 9, we found an age-dependent decreased expression of genes involved in regulation of nervous system processes and synaptic organization and plasticity distinctly in Neil3-deficient AD mice. In contrast, female AD mice presented less differences on the transcriptome level with fewer significant GO processes identified (Fig. 4B). Cluster 1 showed an age-dependent decrease of genes involved in receptor signaling and activity regulation whereas in cluster 5 we found a similar increase in inflammation-related processes as for male AD mice. However, no differences in gene expression upon loss of Neil3 in these clusters in female AD mice were identified. This indicates that Neil3 induces sex-dependent changes in AD on a transcriptional level.

### Hippocampal oxidative DNA damage is increased in Neil3-deficient female mice early in AD

Since *Neil3* is expressed in the hippocampus [52] and oxidative stress-induced DNA damage occurs brain region-specific [8], we examined level of oxidative DNA base lesions in hippocampal tissue of the AD mouse strains. Interestingly, measurement of 5OH-dC (Fig. 5A) and 8oxo-dG (Fig. 5B) showed a significantly higher amount of oxidative DNA damage in female AD mice upon loss of Neil3 at 1 month compared to female APP/PS1 mice. Male Neil3-deficient AD mice showed a similar trend to increased DNA damage level at 1 month, however, the results were not statistically significant. Similar to the DNA damage level in whole brain, we found no accumulation of oxidative base lesions in APP/PS1 mice with age. However, Neil3-deficient AD mice demonstrated a significant decrease in oxidative DNA base lesions from 1 month to 6 months of age (Fig. 5). These data indicate that Neil3 is important for repair of oxidative DNA base lesions or regulation of oxidative stress-induced DNA damage in the hippocampus early in AD pathogenesis.

**Fig. 5.**
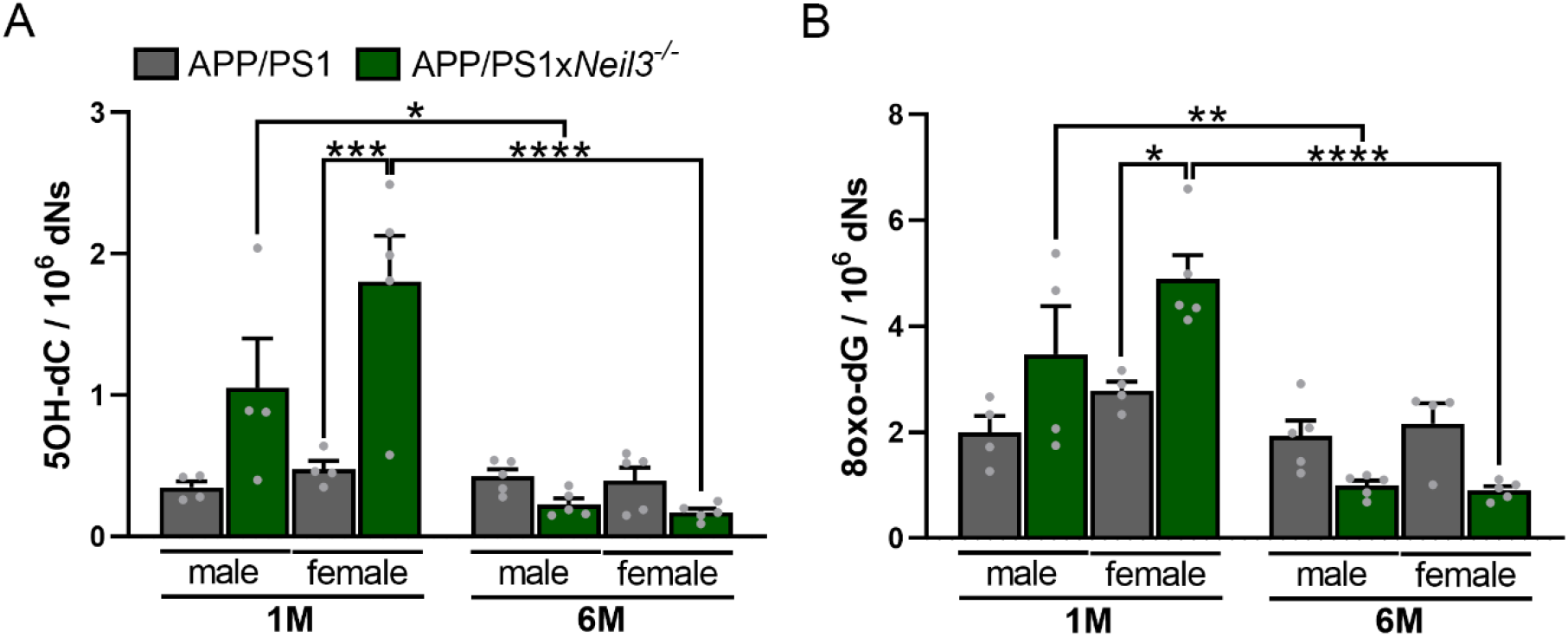
Increased accumulation of oxidative base lesions in the hippocampus of young Neil3-deficient AD mice. (A) Levels of oxidized base lesions 5-hydroxy cytosine (5OH-dC) and (B) 8-oxoguanine (8oxo-dG) in hippocampal samples of APP/PS1 and APP/PS1x*Neil3^−/−^* mice at 1 month (1M) and 6 months (6M) of age measured by using mass spectrometry. Data are shown as mean + SD; n=4-5 per group. *p < 0.05, **p <0.01, *** p<0.001, ****p < 0.0001.

## DISCUSSION

In the present study we generated a novel mouse model deficient for Neil3 to study the role of impaired oxidative base lesion repair in AD pathogenesis. We found that Neil3 altered AD-relevant processes such as amyloidosis, adult neurogenesis, memory and oxidative stress in an age and sex-specific manner. Whereas female Neil3-deficent mice showed significantly less Aβ plaque deposits and increased oxidative DNA damage, male Neil3-deficent mice presented with reduced number of proliferating neural stem cells and impaired working memory during AD progression.

Sex differences in AD are well known but poorly considered when studying AD pathogenesis in animal research [3, 16]. Women have faster cognitive deterioration in MCI and AD [19, 24, 38, 62]. Several studies have shown higher amyloid load in female AD mice [10, 23, 33], which is in line with our findings in the APP/PS1 mice. Interestingly, loss of Neil3 prevented this sex-dependent increase in Aβ accumulation in female mice. Plaque deposition can be altered through estrogen receptor (ER) activation [9, 68, 70]. Notably, DNA glycosylases OGG1 and MUTYH have been shown to affect ER signaling in brain [4] and DNA glycosylases MPG and TDG directly interact with ER to alter transcription [12, 35]. Hence, Neil3 may have a similar function preventing amyloidogenesis. Together, our data suggest that Neil3 affects plaque burden in a sex-dependent manner potentially by altering hormone signaling in AD mice.

Neil3 is a DNA glycosylase known to repair oxidative base lesions. However, previous studies found no differences in levels of oxidative base lesions after myocardial infarction or during prion disease upon loss of Neil3 [28, 49]. Similarly, we found no accumulation of the NEIL3 substrate 5OH-dC in the brain of Neil3-deficient AD mice. This indicates no correlation between Aβ toxicity and oxidative stress in our mouse model, although a delayed increase of oxidative base lesions due to amyloid toxicity at later stages of AD cannot be excluded.

Neil3 was shown to affect learning and memory in aged mice [52]. Here, we find that loss of Neil3 in AD leads to age-dependent impaired working memory in male but not female mice at 6 months of age. At this time point, Neil3-deficient mice in a non-AD background demonstrated similar working memory as their respective controls. The study by Regnell et al. [52] however found impaired memory in 18 months old male *Neil3^−/−^* mice. Thus, our results indicate an age-dependent and AD-specific impact of Neil3 on cognitive function. Gene expression of *Neil3* was previously found most prominent during early stages of postnatal development and to decrease with age [54], thus Neil3 must alter pathways important for memory formation early during AD pathogenesis or in distinct cells of the adult brain. It is believed that amyloid plaques are associated with cognitive decline in AD [36, 37]. We see no correlation of plaque number and memory impairment in our AD model as female AD mice with the highest plaque burden show similar working memory as male AD mice.

Adult neurogenesis is important for cognitive function. In fact, Hollands et al. [25] were the first to demonstrate that reduction of neurogenesis induced memory and learning deficits in a mouse model of AD. Tobin et al. [63] showed that neurogenesis in humans declines with age, and that was associated with cognitive dysfunction in MCI and AD patients. Neil3 is expressed in the neurogenic niches of the brain [22, 55], and has been shown to contribute to neurogenesis after brain injury in postnatal mice [52, 56]. Here, we demonstrate that depletion of Neil3 in AD results in lower number of proliferating stem cells in the hippocampus of male mice indicating that Neil3 contributes *in vivo* to adult hippocampal neurogenesis during AD. This is in line with a previous study that demonstrated prevention of *in vitro* expansion of adult hippocampal neural stem cells upon loss of Neil3 [52]. However, the impact of Neil3 on adult neurogenesis in AD was only evident in young mice as proliferation capacity of neural stem cells was almost completely lost in aged AD mice. It has previously been demonstrated that adult neurogenesis decreases with age and during AD progression [44, 67]. Interestingly, estrogens show a neuroprotective role, increasing neurogenesis in rodent studies [6, 61]. Thus, Neil3-dependent hormonal alterations might contribute to promote neurogenesis in the AD mouse model.

In mouse models of AD, impaired hippocampus-dependent learning and memory was correlated with decreased neurogenesis in the DG [27, 67]. It is tempting to speculate that Neil3 contributes to cognitive function by maintaining adult neurogenesis as we found reduced stem cell proliferation exclusively in Neil3-deficient male AD mice that also presented with impaired working memory. DNA damage accumulation in the hippocampus seems neither contributing to impaired adult neurogenesis nor memory as we found increased level of oxidative base lesion particularly in female AD mice lacking Neil3. Surprisingly, not only did the NEIL3 substrate 5OH-dC, accumulate in Neil3-deficient AD hippocampus but also 8-oxodG, a common marker for oxidative stress. NEIL3 has been shown to excise oxidation products of 8-oxodG but not 8-oxodG itself [71], suggesting that loss of Neil3 might increase oxidative stress in general in the AD mouse model. Moreover, the level of oxidative base lesions in the hippocampus decreased during AD progression in the Neil3-deficient mice. This might be explained by an active removal of cells with a high oxidative stress burden in the hippocampus during AD. This is in line with previous studies showing that excessive DNA damage leads to cell elimination through apoptosis as a protective mechanism [11, 13, 17, 29, 32].

## CONCLUSION

We present for the first time a role of DNA glycosylase Neil3 in AD pathogenesis. We show that Neil3 aggravates amyloid pathology, promotes adult hippocampal neurogenesis, and maintains memory function in a sex-and age-dependent manner but independent of global oxidative base lesion repair in a mouse model of AD. Our findings further confirm that impaired adult neurogenesis is an early event in AD that correlates with memory formation. Moreover, our study underlines the importance of considering sex differences when studying mouse models of AD.

## LIST OF ABBREVIATIONS

Aβ: amyloid beta
Aβ42: amyloid beta peptide 42
AD: Alzheimer’s disease
APP: amyloid precursor protein
BER: base excision repair
DG: dentate gyrus
ER: estrogen receptor
EZM: elevated zero maze
IF: immunofluorescence
IHC: immunohistochemistry
MCI: mild cognitive impairment
MS: mass spectrometry
NEIL3: Nei endonuclease VIII-like 3
NFTs: neurofibrillary tangles
ROS: reactive oxygen species
ssDNA: single-stranded DNA
WT: wild type
5OH-dC: 5-hydroxy-deoxycytosine
8oxo-dG: 8-oxo-deoxyguanosine

## DECLARATIONS

### Ethics approval

All experiments were approved by the Norwegian Animal Research Authority and conducted in accordance with the laws and regulations controlling experimental procedures in live animals in Norway and the European Union’s Directive 2010/63/EU.

### Consent for publication

Not applicable.

### Availability of data and material

The RNA sequencing dataset supporting the conclusions of this article is available in the NCBI’s Gene Expression and Ominbus repository, accession number GSE197199 and https://www.ncbi.nlm.nih.gov/geo/query/acc.cgi?acc=GSE197199.

### Authors’ contributions

M.A.E. and S.S. contributed equally to experiments and analysis, while M.A.E. and K.S. were major contributors in writing the manuscript. A.B.N.N contributed to IHC staining and analysis, T.S. performed the RNA sequency analysis, while N.B. and K.S. conducted behavioral studies. M.B. and K.S. designed and supervised the project. All authors read and approved the final manuscript.

## Acknowledgements

We want to thank the Proteomics and Modomics Experimental Core Facility (PROMEC) at NTNU for the help with mass spectrometry analysis, as well as the Cellular and Molecular Imaging Core Facility (CMIC) at NTNU for the help with scanning mouse brain sections.

