## Supplementary Figures for "Age- and sex-dependent effects of DNA glycosylase Neil3 on amyloid pathology, adult neurogenesis, and memory in a mouse model of Alzheimer’s disease"

**SUPPLEMENTARY INFORMATION**

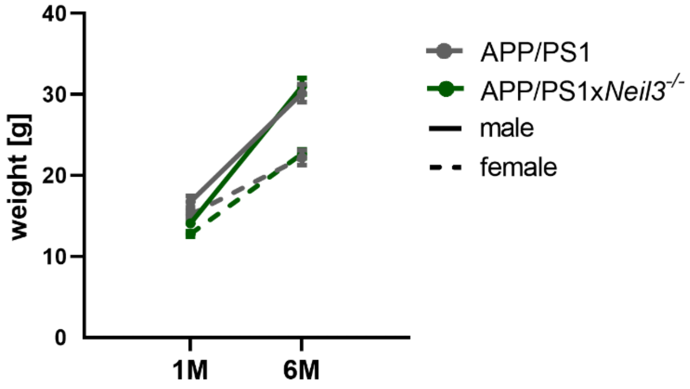


**Additional file 1: Fig. 1 No differences in weight of APP/PS1 and APP/PS1x*Neil3*$^{-/-}$ mice at 1 month (1M) and 6 months (6M) of age**. Data are shown as mean + SD, n=5-9.

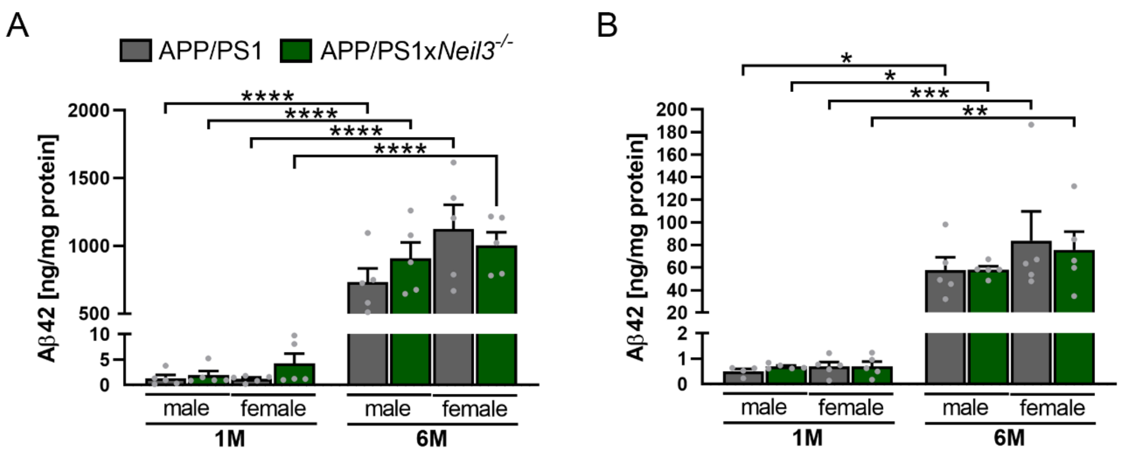


**Additional file 2: Fig 2 No differences in cerebral level of amyloid-β42 (Aβ42) in AD mice upon loss of Neil3.** ELISA analysis of (A) Guanidine-soluble Aβ42 and (B) buffer-soluble Aβ42 in the brain of APP/PS1 and APP/PS1x*Neil3*$^{-/-}$ mice at 1 month (1M) and 6 months (6M) of age. Data are shown as mean + SD; n=5 per group.

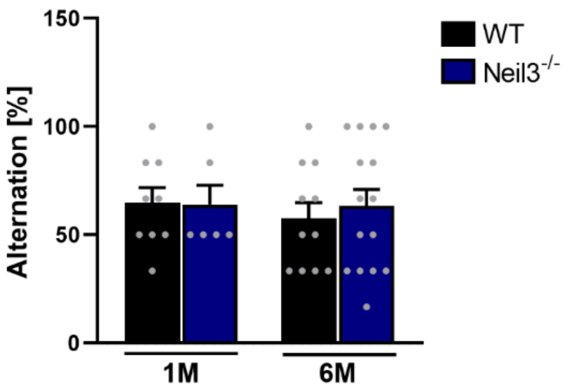


**Additional file 3: Fig. 3 Normal working memory of Neil3-deficient male mice.** Alternation rate of wild type (WT) and *Neil3*$^{-/-}$ male mice assessed by using a T-maze. Data are shown as mean + SD; n=6-15 per group.

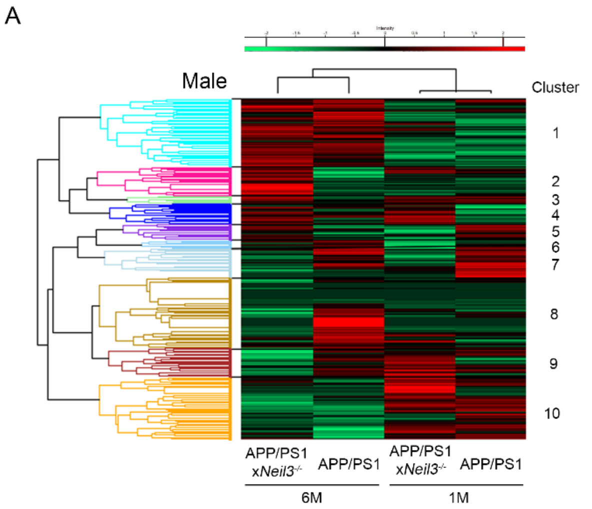


| Cluster | GO biological process | Fold Enrichment | FDR |
|---|---|---|---|
| 1 | antigen processing and presentation of exogenous peptide antigen | 4.2 | 0.012 |
| | ribosomal small subunit assembly | 3.8 | 0.01 |
| | antigen processing and presentation of endogenous peptide antigen | 3.6 | 0.009 |
| | cellular response to interferon-beta | 3.1 | 0.001 |
| | positive regulation of T cell mediated cytotoxicity | 2.7 | 0.02 |
| 2 | oxidative phosphorylation | 3.5 | 0.008 |
| | immunoglobulin production | 3.2 | 5.6E-5 |
| | respiratory electron transport chain | 3.1 | 0.028 |
| 9 | regulation of receptor localization to synapse | 4.9 | 0.05 |
| | synapse organization | 2.22 | 0.001 |
| | regulation of nervous system process | 2.2 | 0.03 |
| | regulation of synapse organization | 2.19 | 0.002 |
| | regulation of synaptic plasticity | 2.17 | 0.01 |
| 10 | regulation of postsynaptic membrane potential | 2.67 | 0.002 |
| | chemical synaptic transmission, postsynaptic | 2.6 | 0.03 |
| | cellular protein complex disassembly | 2.56 | 0.02 |
| | regulation of synaptic vesicle exocytosis | 2.35 | 0.003 |
| | synapse assembly | 2.14 | 0.02 |

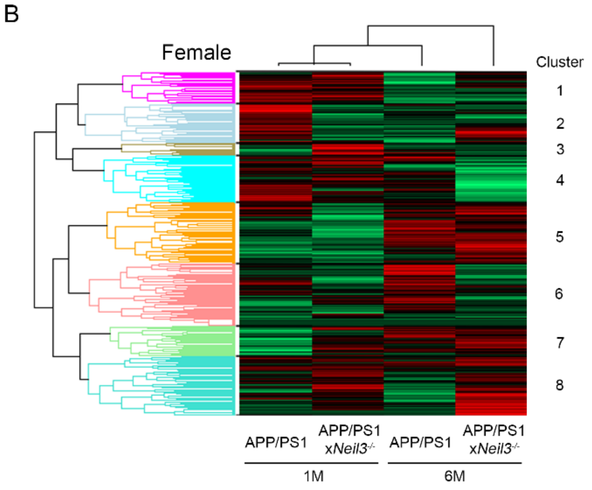


| Cluster | GO biological process | Fold Enrichment | FDR |
|---|---|---|---|
| 1 | regulation of AMPA receptor activity | 3.89 | 0.049 |
| | neuromuscular junction development | 3.49 | 0.003 |
| | glutamate receptor signaling pathway | 3.3 | 0.039 |
| | regulation of calcium ion-dependent exocytosis | 3.1 | 0.044 |
| | synapse assembly | 2.99 | 0.0008 |
| 5 | antigen processing and presentation of endogenous peptide antigen | 4.25 | 0.005 |
| | regulation of CD8-positive, alpha-beta T cell activation | 4.12 | 0.034 |
| | mast cell activation | 3.89 | 0.004 |
| | antigen processing and presentation of exogenous peptide antigen | 3.46 | 0.02 |
| | cellular response to interferon-beta | 3.2 | 0.004 |

**Additional file 4: Fig. 4 RNA sequencing reveals differences in nervous system processes with age and upon loss of Neil3 in male AD mice.** Heatmap with hierarchical clustering and associated GO biological processes in brain of (A) male and (B) female APP/PS1 and APP/PS1x*Neil3*$^{-/-}$ mice at 1 month (1M) and 6 months (6M) of age.
